# sgDI-tector: defective interfering viral genome bioinformatics for detection of coronavirus subgenomic RNAs

**DOI:** 10.1101/2021.11.30.470527

**Authors:** Andrea Di Gioacchino, Rachel Legendre, Yannis Rahou, Valérie Najburg, Pierre Charneau, Benjamin D Greenbaum, Frédéric Tangy, Sylvie van der Werf, Simona Cocco, Anastasia V Komarova

## Abstract

Coronavirus RNA-dependent RNA polymerases produce subgenomic RNAs (sgRNAs) that encode viral structural and accessory proteins. User-friendly bioinformatic tools to detect and quantify sgRNA production are urgently needed to study the growing number of next-generation sequencing (NGS) data of SARS-CoV-2. We introduced sgDI-tector to identify and quantify sgRNA in SARS-CoV-2 NGS data. sgDI-tector allowed detection of sgRNA without initial knowledge of the transcription-regulatory sequences. We produced NGS data and successfully detected the nested set of sgRNAs with the ranking M*>*ORF3a*>*N*>*ORF6*>*ORF7a*>*ORF8*>*S*>*E*>*ORF7b. We also compared the level of sgRNA production with other types of viral RNA products such as defective interfering viral genomes.

## Introduction

Viral RNA-dependent RNA polymerases (RdRp) ensure multiple molecular mechanisms to produce a large spectrum of viral RNA products inside infected cells. Some of these molecular mechanisms have already been described in detail and others are yet to be uncovered. Mechanisms of RNA virus replication and transcription, cap-snatching, and RNA editing have been relatively well described [Strauss and Strauss, 2007], whereas molecular mechanisms underlying defective viral genome (DVG) production have yet to be discovered. DVGs are truncated forms of and/or rearranged viral genomes generated by most viruses during viral replication. DVG can be also called defective interfering (DI) genomes when viral particles containing them are able to interfere with standard virus replication [Pathak and Nagy, 2009]. DVG and viral genomes share the minimum essential characteristics for replication: a competent initiation site at the 3’-end and its complementary sequence at the 5’-end. However, DVG genomes are defective for replication in the absence of the complete functional virus genome that provides the missing functions. Four main classes of DVGs exist: deletions, insertions, snap-back DI genomes or “hairpin” structures, and copy-back or “panhandle” structure DI genomes (see Fig. 1; [Lazzarini et al., 1981, Dimmock et al., 2014]).

**Figure 1:**
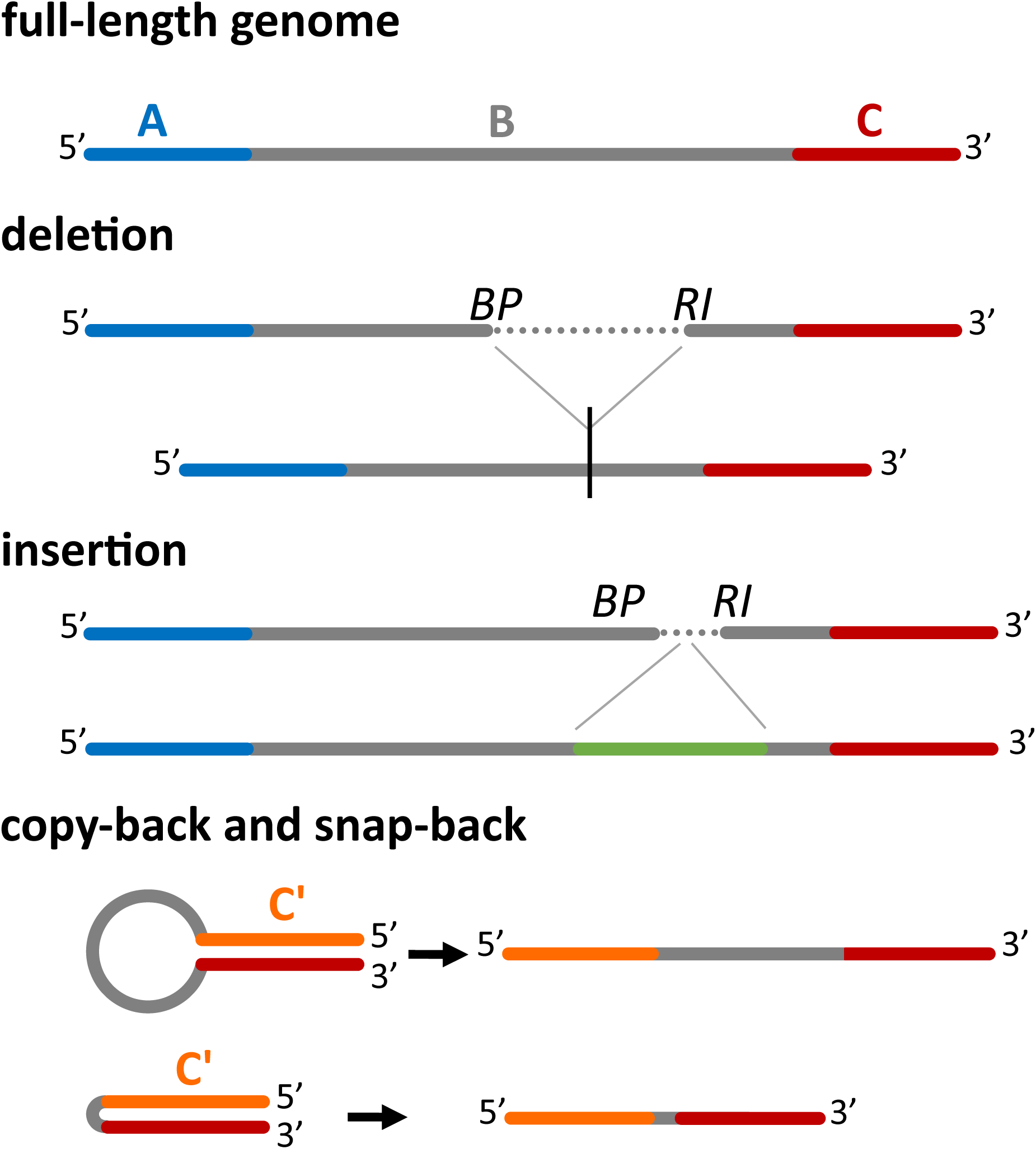
Four main classes of DI genomes can be detected by DI-tector. The full-length genome (divided here in three regions, A, B and C) is shown first. When a part of the region B is missed then DI-tector detects a deletion, if instead a region is added, DI-tector detects an insertion. Copy-backs and snap-backs are formed through junctions involving the two strands (positive-sense and negative-sense). C’ is the region complementary to C. BP: breakpoint site, RI: reinitiation site.

SARS-CoV-2 is an enveloped, positive-sense, single-stranded RNA virus with a genome of nearly 30,000 nucleotides (nts) [Lu et al., 2020]. Similar to other coronaviruses, SARS-CoV-2 replication involves the synthesis by the viral RdRp of positive and negative sense full-length genomes as well as the production of a nested set of subgenomic RNA (sgRNAs). SARS-CoV-2 sgRNAs encode four structural proteins (S, spike; E, envelope; M, membrane; N, nucleocapsid) and several accessory factors (ORF3a, ORF3b, ORF6, ORF7a, ORF7b, ORF8, and ORF10) [Davidson et al., 2020, Kim et al., 2020, Wu et al., 2020, Zhou et al., 2020].

sgRNAs are produced by the yet to be determined complex mechanisms assigned as discontinuous transcription that includes two steps [Pasternak et al., 2006]. The first consists in the viral RdRp pausing the negative-sense RNA synthesis at specific 6-7 nt in length sequence (body transcription regulatory sequence, TRS-B) at the 3’ of the viral genome and then performing a long-range jump at 5’ end of the genome to join a common sequence of approximately 70 nts encompassing another, identical 6-7 nt in length sequence. This second hexanucleotide sequence is named leader transcription-regulatory sequence (TRS-L), and the part of the viral genome starting at the 5’-end and ending just after the TRS-L is called leader sequence. In our study we use generically TRS to indicate short sequences which are found both in the final part of the leader sequence, and in the 3’ part of the genome around the site of the junction. The second step is the replication of the positive sense sgRNA from the negative-sense RNA template produced at the first step. This way, the molecular organization of coronavirus sgRNAs is similar to the deletion type of the DI genomes, and methods applied for DI genome detection in NGS data become suitable for the detection of coronavirus sgRNA. However, the subgenomic biogenesis mechanism is still under intensive study, and several important questions have been addressed only very recently: for instance, it has been suggested that in addition to the full-length SARS-CoV-2 genomic template the sgRNAs themselves can give rise to shorter sgRNA, through additional RdRp pause-and-jump events [Wang et al., 2021]. Another question that has been addressed recently, with a particular focus on SARS-CoV-2, regards the role in sgRNA expression of the secondary structure within the 5’ untranslated region [Sola et al., 2015, Miao et al., 2021].

Several sgRNA-oriented NGS studies have already provided various ratios of the nested SARS-CoV-2 sgRNA concentrations. Kim *et al*.’s study applied nanopore direct RNA sequencing validated by DNA nanoball sequencing (MGI NGS platform) to demonstrate that Vero cells infection with SARS-CoV-2 produces a complexed transcriptome with 9 canonical sgRNAs and non-canonical viral ORFs [Kim et al., 2020]. To detect sgRNA reads from short length NGS data Kim *et al*. applied Spliced Transcripts Alignment to a Reference (STAR) free open source software [Dobin et al., 2012], and performed a quantitative analysis of sgRNA transcription. They revealed that N RNA was the most abundantly expressed transcript, followed by S, 7a, 3a 8, M, E, 6, and 7b RNAs [Kim et al., 2020]. Finkel *et al*. used two approaches to calculate the abundance of sgRNAs in NGS data from SARS-CoV-2 infected cell cultures [Finkel et al., 2021]. The first was the STAR-based assessment of the relative abundances of RNA reads spanning leader–body junctions for the canonical sgRNAs. The second approach used deconvolution of RNA densities. In the deconvolution or “decumulation” approach the RNA expression of each ORF is calculated by subtracting the RNA-read density upstream of the ORF region (inter-TRS region) [Irigoyen et al., 2016]. Finkel *et al*.’s study has described N transcript as the most abundant followed by M, ORF7a and ORF3a. Finally, the first bioinformatic pipeline for detection and quantification of sgRNA specifically in SARS-CoV-2 genomic sequencing data has been proposed and called Periscope [Parker et al., 2021]. Periscope deals with various types of SARS-CoV-2 sequencing analysis including ARTIC Nanopore-generated and Illumina metagenomic sequencing. In order to identify sgRNAs, Periscope requires previous knowledge of the coronavirus leader sequence [Parker et al., 2021]. However, the above approaches are rather complex, requiring several intermediate steps or previous experience with a number of other bioinformatic tools, especially when applied to SARS-CoV-2 short-read NGS data to detect and characterize sgRNAs.

We have recently developed a user-friendly bioinformatic pipeline called DI-tector to detect various types of DVGs from NGS data [Beauclair et al., 2018]. In this report, we extend the functionality of DI-tector by introducing sgDI-tector, a tool that, after running DI-tector, can properly detect and quantify coronavirus sgRNAs without previous knowledge of the TRS. We use the output of sgDI-tector to compute the SARS-CoV-2 sgRNA ratios and discuss the comparison with the results obtained through previously developed approaches [Kim et al., 2020, Finkel et al., 2021, Parker et al., 2021], showing its largest robustness and sensitivity. To confirm sgDI-tector’s potential to detect rare non-canonical sgRNA populations from NGS data RT-qPCR approach was applied. Moreover contrary to other methods, sgDI-tector does not impose a junction sequence, allowing us to investigate the sequences found at the junction and their variability.

## Results

### DI-tector detects various types of SARS-CoV-2 DI genomes

First, we generated an NGS data set on total RNA from human cells (HEK293 transduced with ACE) infected with SARS-CoV-2. Then we ran DI-tector on the RNAseq data to characterize and quantify DVGs. The results of RNAseq and alignment to reference genomes, and those of DI-tector, are presented in Fig. 2 (the raw number of counts used for this Figure are provided in Suppl. Table 1). We observed a relevant number of reads mapped to SARS-CoV-2 genome (respectively 33%, 40%, and 40% in the 3 biological replicates, see Suppl. Table 1), and a much smaller quantity of DVG reads. The most represented type of DVG read was deletions, which accounted for about 74% of the DVG reads (Table 1). While expected, given the coronaviruses’ mechanism of production of sgRNA, this large fraction of deletions suggests that DI-tector results can be used to identify and quantify viral sgRNA from NGS data, in a simple and efficient way. Indeed, we were able to associate 68% of these DVG reads to canonical SARS-CoV-2 sgRNA transcripts (coding for ORFs: S, 3A, E, M, 6, 7a, 7b, N), see Material and Methods section for details about the pipeline used. We also found a quite large number of insertions, accounting for about 24% of the total DVG reads. Finally, we observed around 2% of the total DVG of the copyback or snap-back type of DI genomes in SARS-CoV-2 infected cells (Table 1, Suppl. Table 1).

**Figure 2:**
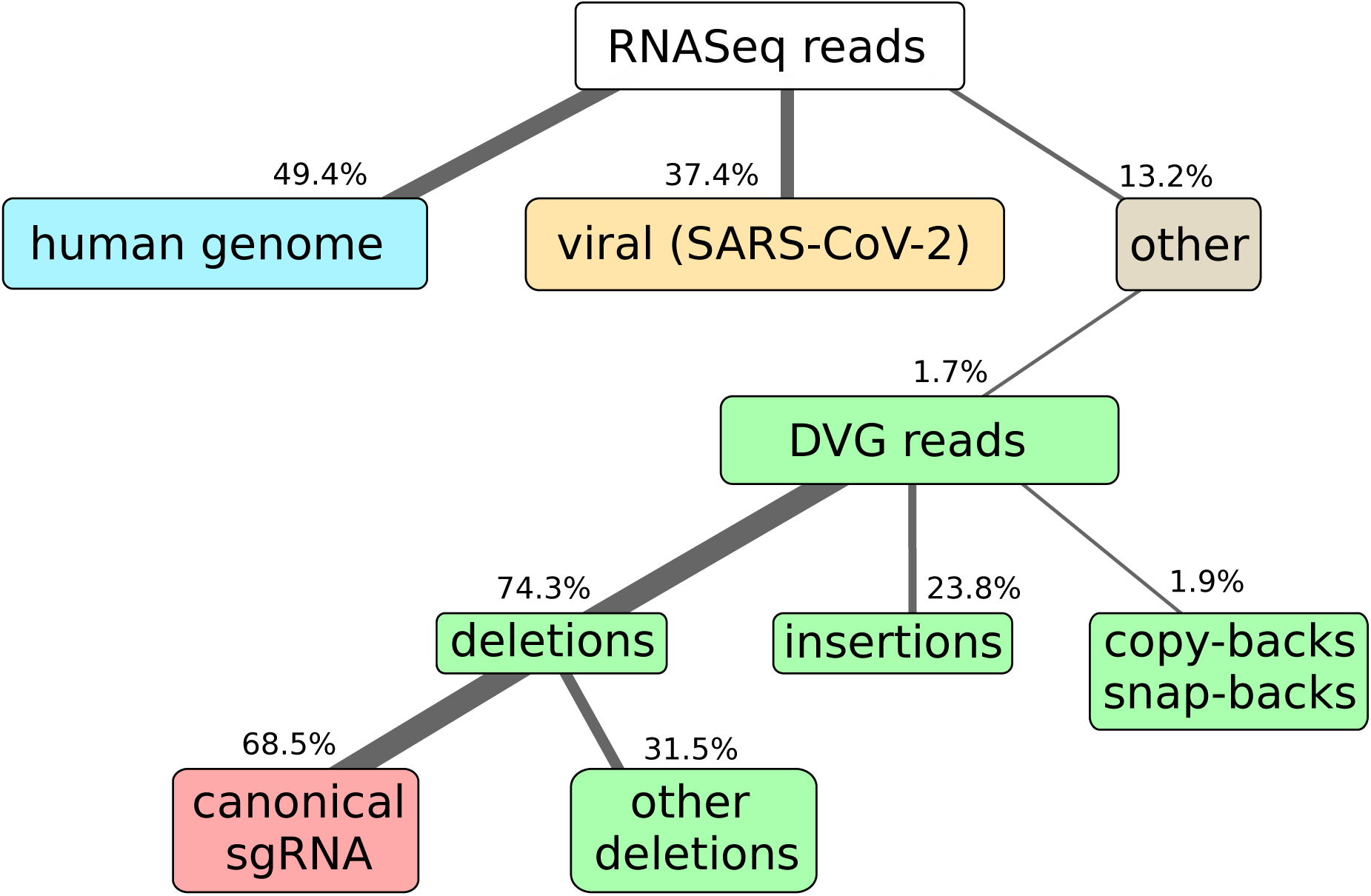
Most of the DVG reads can be associated to canonical sgRNAs. Here we show the results of RNAseq and alignment of the reads to the human and SARS-CoV-2 genomes. NGS library preparation was performed with an ribodepletion step. The unmapped reads have been further processed with DI-tector and the resulting characterization of the DVG reads into deletions, insertions and copy-backs/snap-backs is given. The percentage of junction reads corresponding to canonical sgRNA, that is standard annotated subgenomic ORFs for SARS-CoV-2 (S, 3A, E, M, 6, 7a, 7b, N, 10), is specified. All percentages are averaged over three biological replicates. Cells colors are given to classify reads in: host-related reads (blue), viral reads (yellow), DVG reads (green), canonical sgRNA reads (red), other reads (gray).

**Table 1:**
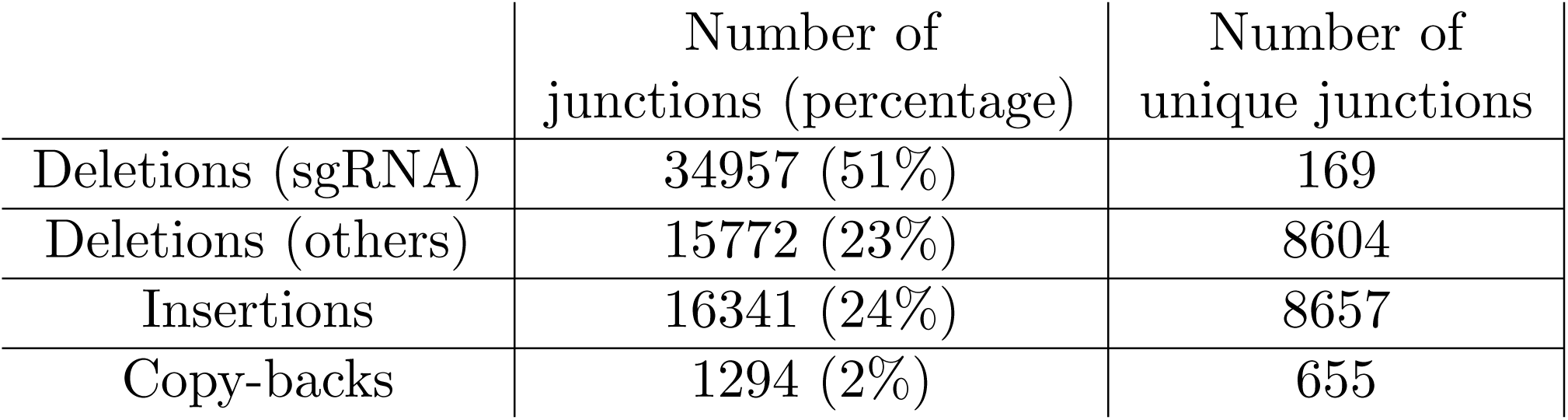
Overview on DVG observed in this study. All the values are obtained as averages of 3 biological replicates. Unique junctions are defined as different BP-RI positions spanned by observed reads. Copy-back DVGs also include snap-back DVGs.

|  | Number of<br>junctions (percentage) | Number of<br>unique junctions |
| --- | --- | --- |
| Deletions (sgRNA) | 34957 (51%) | 169 |
| Deletions (others) | 15772 (23%) | 8604 |
| Insertions | 16341 (24%) | 8657 |
| Copy-backs | 1294 (2%) | 655 |

### DVGs of deletion type can be used to characterize the nested set of sgRNAs

The leader-body junctions formed during the transcription of sgRNA in coronaviruses are detected by DI-tector as deletion DVGs, and the importance of such sgRNAs is reflected by their abundance with respect to other DVG types. We assessed here the possibility of using DI-tector to characterize the nested set of ORFs transcribed as sgRNAs, and to compare the levels of expression of different sgRNAs. Our pipeline, which we named sgDI-tector, starts from the DI-tector output and is based on two assumptions: (i) sgRNA coding for expressed ORFs are transcribed more frequently (on the overall) than classical DVGs; and (ii) there is a leader sequence shared among all sgRNAs. The detailed pipeline is described in the Method section. We stress, however, that differently from other methods we do not need to explicitly know the leader/junction TRS to apply our algorithm, but the fact that there is one is necessary for the algorithm to work. We run our analysis by using three biological replicates, and all of the results presented here are almost unchanged in each of them (Fig. 3). As can be seen in Fig. 3, in each of our samples we observed clear signals of several ORFs, in particular M, 3a, N, 6, 7a, 8, S and E. We also found some signal of direct transcription of ORF 7b, which could also be translated from the same sgRNA coding for ORF 7a as suggested for SARS-CoV in [Schaecher et al., 2007]. Moreover, we observed a leader-body junction resulting in a potential expression of a non-canonical ORF. Remarkably, its number of counts was comparable with ORF E and larger than ORF 7b when counts were averaged over replicates, see Fig. 4. This ORF, which is referred to as “U6cb1” in Fig. 3 and Fig. 4 (in Suppl. Table 2 we reported the AUG codon position in our reference SARS-CoV-2 genome, for each non-canonical sgRNA we detected), codes for a 20-amino-acid long protein and has been previously identified in Finkel *et al*. [Finkel et al., 2021] and referred to as 7b.iORF1. Additional leader-body junctions in our data gave rise to different ORFs which were found in all three biological replicates. Those found with highest count numbers are presented in Fig. 3 and Fig. 4 (blue bars). To provide a control for false positives using sgDI-tector we analyzed as input the RNASeq data coming from mock-infected HEK293-ACE2 cells. sgDI-tector did not give any sgRNA read in this case, confirming the robustness and sensitivity of our approach.

**Figure 3:**
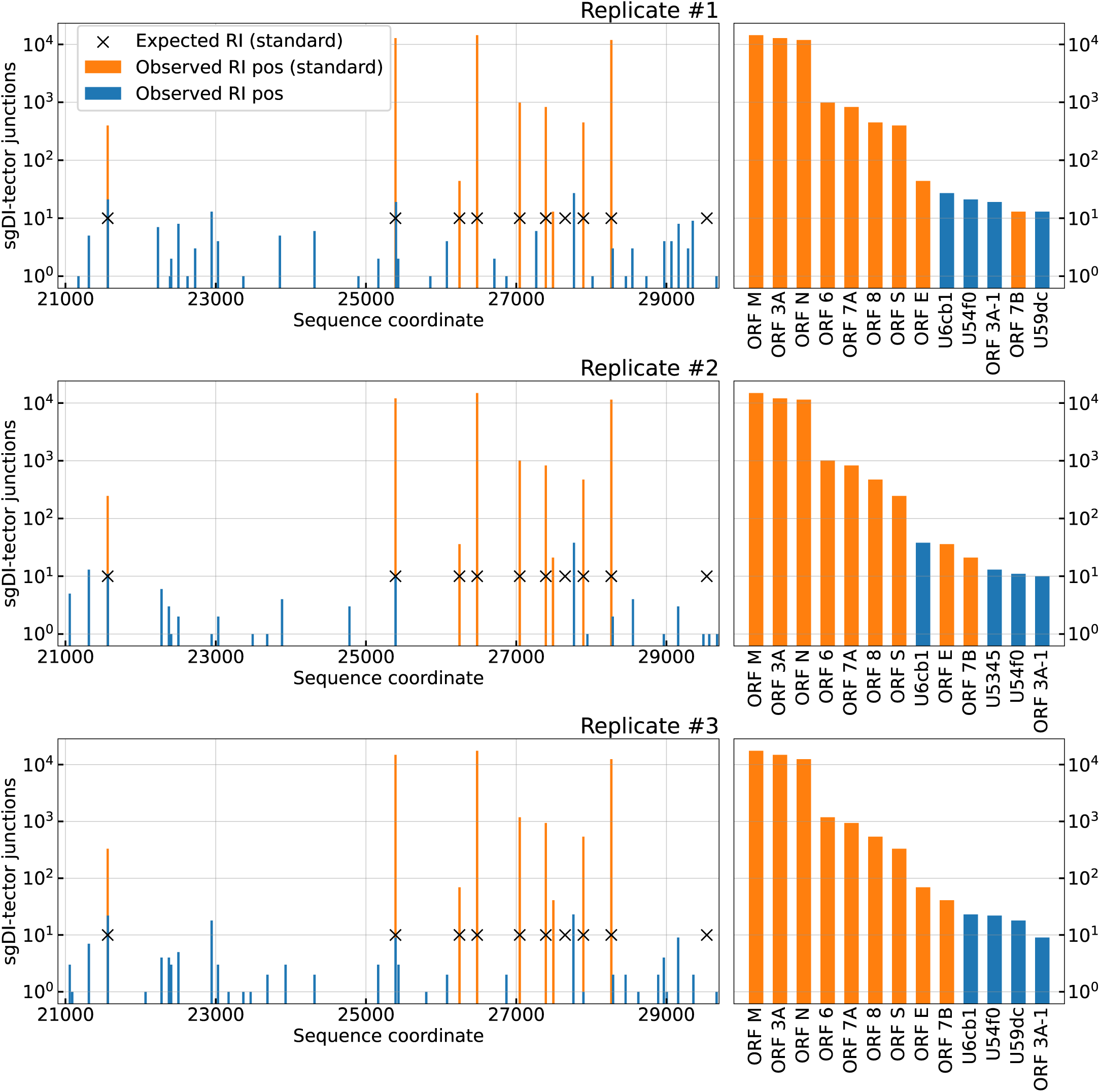
sgDI-tector detects most canonical sgRNAs in all replicates, with high number of counts. Left panels: deletion DVGs distribution across the last 10kb positions of the SARS-CoV-2 genome (GISAID ID: EPI ISL 414631). Each blue or orange bar corresponds to a deletion, the position of the bar being the starting point of the “body” part of the junction (called RI position in sgDI-tector). Crosses are expected RI positions from [Alexandersen et al., 2020] and bars are colored in orange if the deletion is observed in that position in our data. Right panels: number of counts and ORF name for the 13 deletions with most counts observed in each replicate. Orange bars correspond to orange crosses in the left panel and represent canonical TRS. Blue bars correspond to putative non-canonical ORFs detected in our data. Names for non-canonical ORFs are hexadecimal numbers representing the position of the corresponding start codon (AUG) in the standard 5’-to-3’ sense in the reference sequence (GISAID ID: EPI ISL 414631), see also Suppl. Table 2.

**Figure 4:**
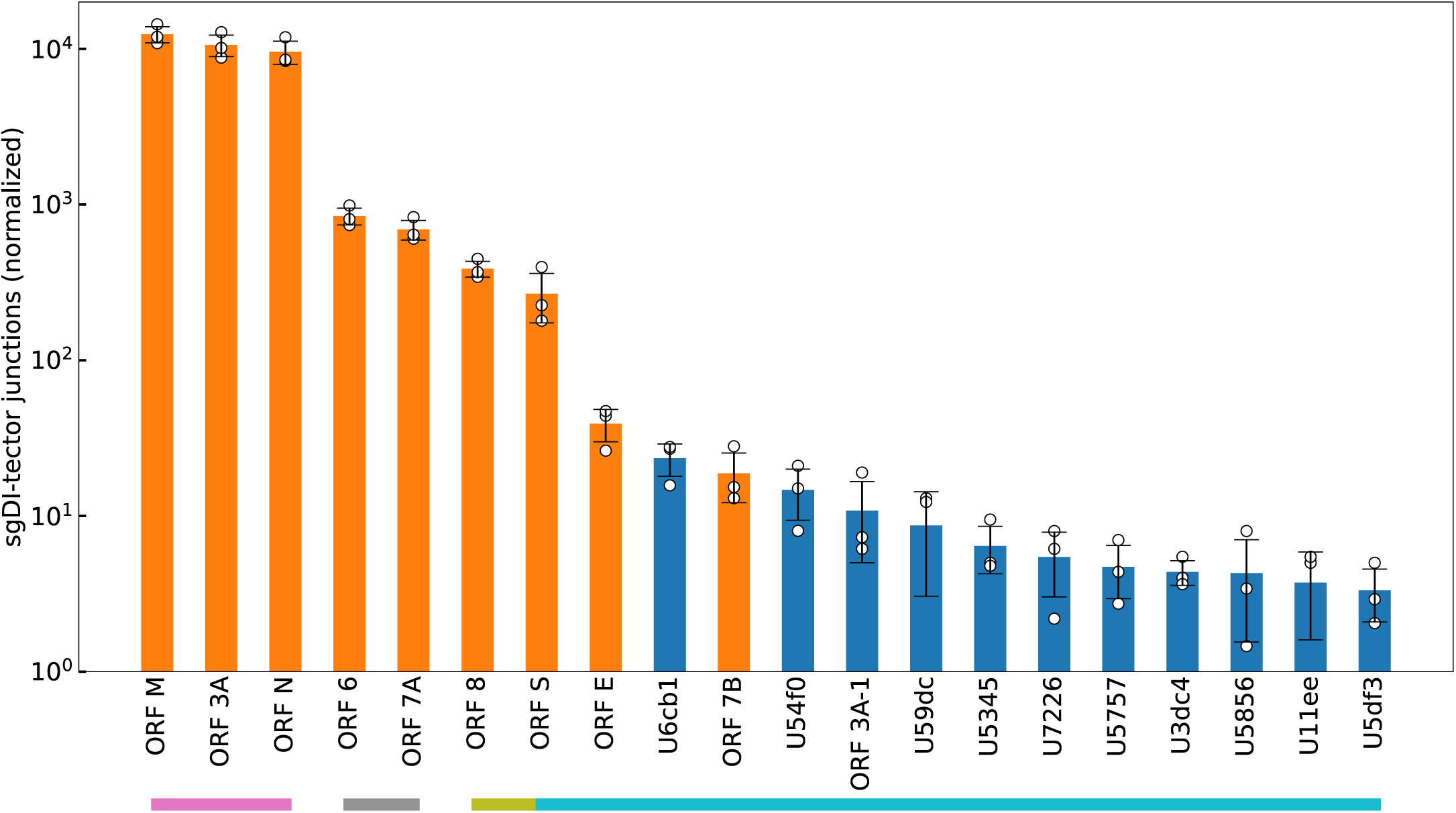
Canonical sgRNAs and some non-canonical sgRNAs are consistently observed across the three biological replicates. ORF names and numbers of counts of the 20 deletions with most counts observed in NGS data in three biological replicates. Data coming from different replicates have been normalized so that the number of viral reads observed in each replicate is constant (see Materials and Methods). Bars heights are given by the average of the three replicates (shown as white dots after normalization), and error bars represent the standard deviation. The colored bars below the ORF names indicate statistical significance of the count differences: ORFs above bars of different colors have statistically different junction counts (p-value *<* 0.05, from a two-sample, two-tailed, Welch’s unequal-variance t-test).

### Validation of non-canonical sgRNA produced inside ORF1ab

We used RT-qPCR approach in order to test accuracy of our sgDI-detector algorithm. We validated the non-canonical ORF U3dc4 that has its body TRS sequence located in ORF1ab. ORF U3dc4 was less present in our data than the majority of other non-canonical ORFs. Of note, U3dc4 was previously detected in [Kim et al., 2020] study by STAR approach, but to our knowledge never validated by a conventional approach. We performed RT-qPCR analysis on total RNA extracted from HEK293 (transduced with ACE) infected with SARS-CoV-2 or mock-infected (negative control) cells. As expected, ORF U3dc4 was detected only in RNA extracted from infected cells similar to the detection of canonical ORF encoding the N protein (Table 2, Suppl. Table 3). These experiments validate the presence of ORF U3dc4 in SARS-CoV-2-infected cells by a RT-qPCR approach and thus highlight the power of sgDI-tector to reveal the landscape of the sgRNA population from NGS data.

**Table 2:** RT-qPCR validation of U3dc4 sgRNA transcript. RT-qPCR *C_t_* values for U3dc4, ORF N, and GAPDH detection in cDNA equivalent of 50 ng of total RNA extracted from SARS-CoV-2 or mock-infected HEK293T cells are shown. Samples were analyzed in duplicates. ND: Not Determined (*C_t_ >* 34). Data are given as average *±* standard deviation. Results obtained from the other biological replicates are given in Suppl. Table 4.

| Target detected | SARS-CoV-2 | Non-infected |
| --- | --- | --- |
| U3dc4 | $27.0 \pm 0.1$ | ND |
| ORF N | $17.0 \pm 0.1$ | ND |
| GAPDH | $20.65 \pm 0.05$ | $20.6 \pm 0.1$ |

### Comparison of sgDI-tector results on our data with existing bioinformatic tools

Several techniques have been used so far to detect sgRNA expression levels in coronaviruses. Firstly, the reads per kilobase of transcript per million mapped reads (RPKM) have been directly used for several coronaviruses, once the RPKM of each sgRNA is properly “decumulated” from downstream ORFs, which are present in each ORF transcript [Irigoyen et al., 2016]. Despite its simplicity, we show in Fig. 5a that, in our case, the decumulation approach gave results which failed to correlate with those obtained with the junction analysis performed by sgDI-tector. Moreover, the decumulation resulted in several ORFs having a negative number of counts, and this can happen for two reasons: the (unavoidable) errors in the estimation of transcription levels from the NGS data (especially for short ORFs) and the presence of other DVGs which are not considered in the decumulation procedure. Similar results have been independently reported in [Finkel et al., 2021]. We believe that the absence of correlation between junction counts and decumulated RPKM in our sample cannot be explained by a failure of sgDI-tector and/or of our pipeline. Indeed we recovered, with the same pipeline, a much higher correlation between junction counts and decumulated RPKM in another NGS dataset, collected by Finkel *et al*. ([Finkel et al., 2021], see Suppl. Fig. 1). In Fig. 5b and Fig. 5c we compare sgDI-tector with other tools which exploit chimeric reads to search for putative sgRNA junctions: Periscope [Parker et al., 2021] and STAR 2.7.3a as used in [Kim et al., 2020, Finkel et al., 2021, Wang et al., 2021]. The most abundant sgRNAs (M, 3A and N) are detected by all tools with many counts, although the ranking of them is different in the output of STAR. Periscope, on the other hand, cannot detect at all ORF E, and ORF 7b (ORF 6 is detected with only one count), suggesting that our tool might be more accurate for Illumina data than Periscope, which has been developed to deal with ARTIC Nanopore-generated sequencing data. DI-tector and STAR results are almost perfectly correlated for most of the sgRNAs. However, junctions associated with ORF 8, and most of the junctions associated with ORF 3A, were detected by STAR only when NGS reads mapped on the negative-sense viral genome were included in the data analysis, thus after performing manual curation of the data. The only ORF for which there is a sensible difference in sgDI-tector’s and STAR’s results is ORF 3A, as STAR can detect significantly fewer junctions with respect to sgDI-tector (see Table 3). Although it is difficult to clearly assess whether the error is on the STAR or sgDI-tector side, the former hypothesis seems more supported since the junction counts of ORF 3A obtained by STAR is statistically different from both the counts obtained by sgDI-tector and Periscope, while the junction counts detected by sgDI-tector is compatible with the result obtained from Periscope. In Table 3 we present a more systematic comparison among the three tools based on the abundance ranking of the canonical sgRNA, whose outcome is that in the only case where the results of sgDI-tector and STAR were not compatible, Periscope’s result was compatible with sgDI-tector’s result and not with STAR’s result. In Fig. 5d-f, we compared our results with those obtained through STAR for two other RNA-sequencing data sets, [Kim et al., 2020, Finkel et al., 2021]. In both publications the authors infected Vero cells, but with different protocols: Finkel *et al*. used a multiplicity of infection (MOI) of 0.2, harvested cells after 5 hours post-infection (hpi) and 24 hpi, and the sequencing was conducted on the Illumina Miseq platform; Kim *et al*. used a MOI of 0.05, harvested cells 24 hpi, and used nanopore and nanoballs RNA sequencing. It is apparent that ORF N is consistently one of the most transcribed sgRNAs. However, several variations between experiments can be noticed: for instance, ORF E seems to be more transcribed in Finkel *et al*.’s analysis [Finkel et al., 2021], and ORF S is the second most-transcribed sgRNA in Kim *et al*.’s analysis [Kim et al., 2020]. In addition to these differences, it is apparent that for most of the ORFs HEK293 cells present less junctions than those observed in previous experiments (see dashed lines in Fig. 5d-f) performed on Vero cells. Differences in efficiency of infection of Vero cell line, that is routinely used to amplify SARS-CoV-2, and of ACE-transduced HEK293 (ST-CH^ACE-2^) cells could explain the observed inequality in number of junctions.

**Figure 5:**
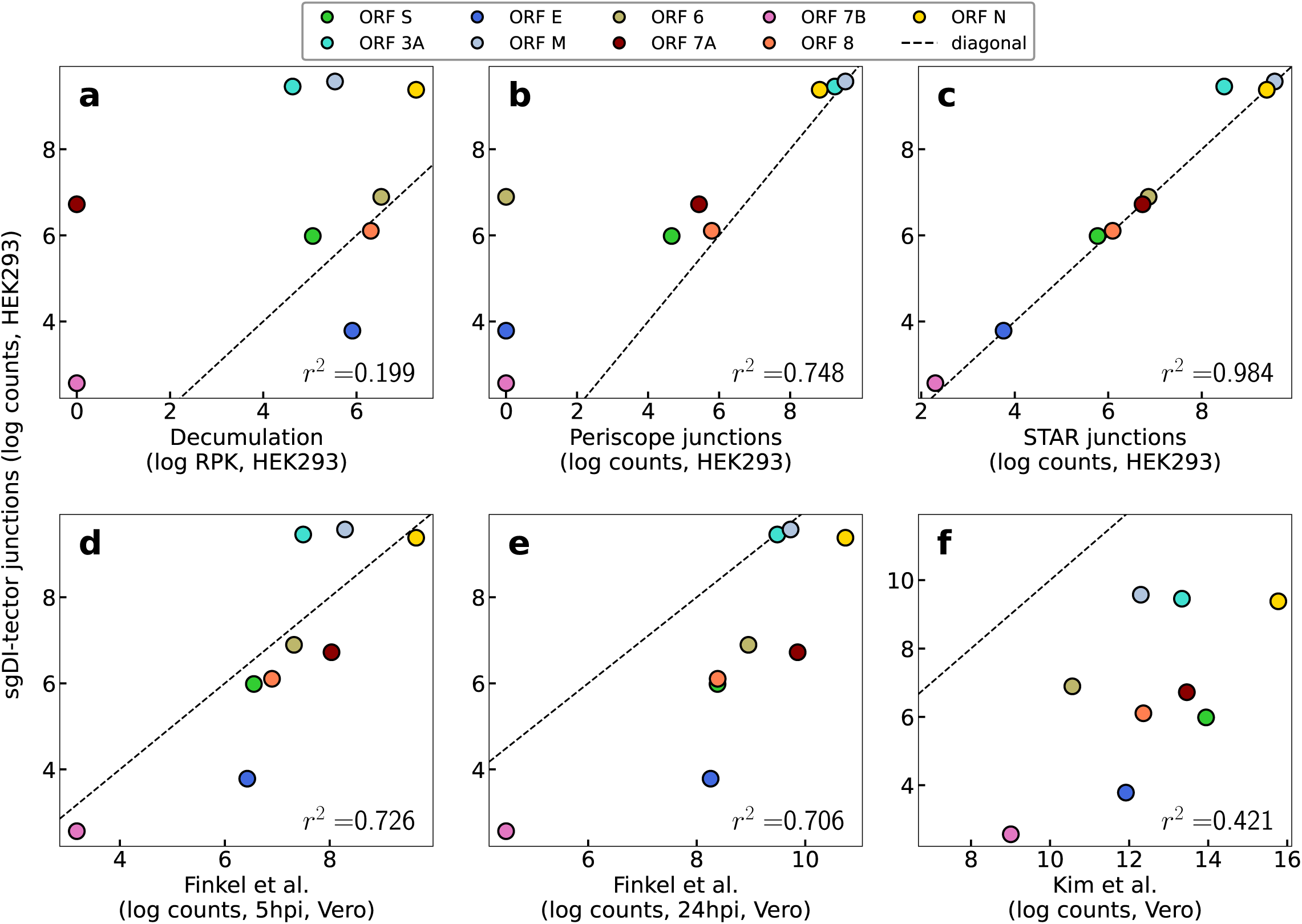
sgDI-tector results are not correlated with decumulation results, while agreeing with other tools applied on the same and on other data. The first row (a-c) presents only results obtained in our experiments and analyzed with several bioinformatic tools, while the second row (d-f) presents a comparison between our data and other data present in the literature. All given results are for one replicate. 1 pseudocount has been added when necessary to visualize the same number of ORFs in each plot. The black dashed line is the diagonal line, added to ease the comparison between the different methods. Notice that the plots on the bottom row compare results on different cell lines: HEK293 on the y axis, and Vero on the x axis.

**Table 3:** Comparison of the sgRNA abundance obtained by sgDI-tector, STAR and Periscope. We compare the average and standard deviation of the counts observed by sgDI-tector, STAR and Periscope for each canonical sgRNA junction (given as mean *±* standard deviation from our 3 biological replicates). A color code is used to indicate the statistical significance as follows: a cell has a green background if it is not statistically different from at least another tool, an orange background if it is statistically different from the other tools, and a red background if no junctions are found in the 3 replicates. ORF 10 is not present as none of the three tools detected the corresponding sgRNA junctions. The p-values used to assign colors are obtained through two-sample, two-tailed, Welch’s unequal-variance t-tests. Counts from different replicates have been normalized so that the number of SARS-CoV-2 reads observed in each replicate is constant (see Materials and Methods).

|  | STAR | sgDI-tector | Periscope |
| --- | --- | --- | --- |
| ORF S | 220 ± 74 | 267 ± 94 | 85 ± 16 |
| ORF 3A | 4239 ± 624 | 10573 ± 1665 | 8663 ± 1375 |
| ORF E | 41 ± 7 | 39 ± 9 | 0 ± 0 |
| ORF M | 12175 ± 1461 | 12396 ± 1466 | 12207 ± 1433 |
| ORF 6 | 817 ± 103 | 843 ± 104 | 0.6 ± 0.4 |
| ORF 7A | 695 ± 104 | 691 ± 99 | 188 ± 35 |
| ORF 7B | 16 ± 7 | 19 ± 7 | 0 ± 0 |
| ORF 8 | 382 ± 46 | 387 ± 45 | 290 ± 31 |
| ORF N | 9633 ± 1640 | 9578 ± 1619 | 5767 ± 806 |

### Applying sgDI-tector on previously published SARS-CoV-2 NGS data sets

Next, we run our tool starting from the raw RNA-seq data obtained by Finkel *et al*. [Finkel et al., 2021] to make a comparison with their results, which is presented in Fig. 6. We observed that the results obtained by sgDI-tector and STAR 2.7.3a are well correlated, showing the effectiveness of the approach proposed here to quantify sgRNA transcription from NGS data. The results of the same test done with Periscope showed a much lower correlation with the original Finkel *et al*.’s junction counts. In particular, Periscope completely missed junctions with sgRNAs 6, E, and 7b (they are shown with 1 pseudocount in Fig. 6c-d) at 5 hpi, which on the opposite were found with a striking correlation by sgDI-tector and by Finkel *et al*.

**Figure 6:**
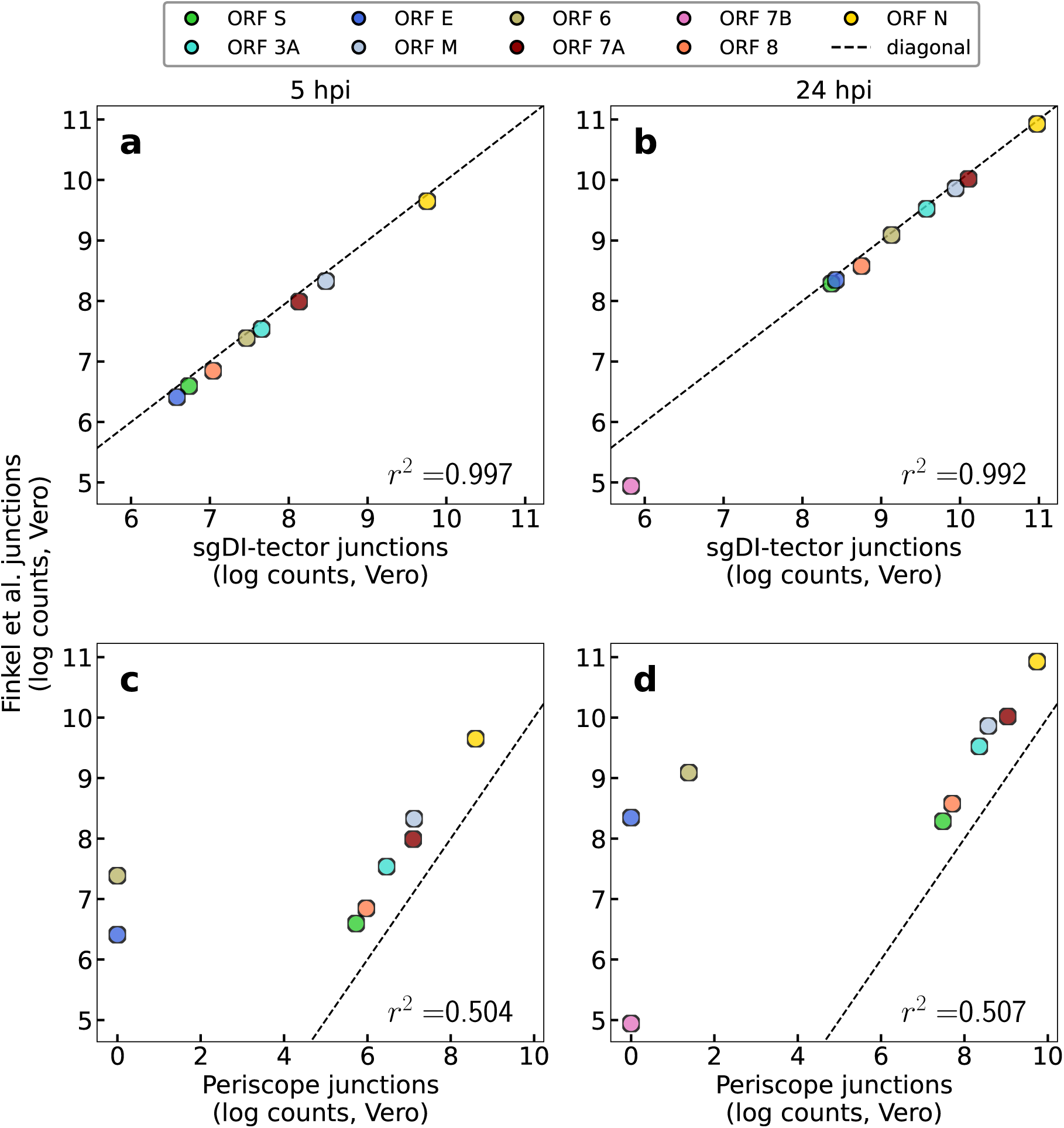
sgRNA junction counts obtained with sgDI-tector correlate with Finkel *et al*.’s junction counts (STAR) while Periscope results show a lower correlation. Left (right) column contains the results for data at 5 (24) hpi. Only data for the first biological replicate are presented here. ORF 10 junctions are never found by both STAR and by DI-tector, while a single read has been detected by Periscope at 5 hpi. 1 pseudocount has been added to junctions which are not detected by one tool while being detected by the other. The black dashed line is the diagonal line, added to ease the comparison between the different methods.

### Leader-body conserved TRSs can be obtained from sgDI-tector output data

sgDI-tector does not require knowing a priori the identity of the leader sequence, as the observed junction-spanning reads are used in the pipeline to recover its position. For the viral strain used in our experiment, the leader-body junctions start, from the 5’ (leader) side, between position 60 and 80. This subsequence can be interpreted as the final part of the leader sequence. Focusing on the nucleotides around the RI positions for the reads detected by DI-tector, our tool allows for a completely automatic discovery of the leader-body conserved TRSs, as shown in Table 4, Fig. 7, Suppl. Table 3 and Suppl. Fig. 2. As apparent from the table and from the logo, the previously reported TRS 5’-ACGAAC-3’ [Wang et al., 2021] has clearly a special role, being present in most cases (and in the junctions with the highest number of observed counts). In particular, the nucleotides AAC in positions 71-72-73 (last part of 5’-ACGAAC-3’) are perfectly conserved within the body partners of the leader-body junctions, for all the junctions analyzed here. Remarkably, however not all the junctions present the 5’-ACGAAC-3’ motif. Moreover, this analysis shows that all the putative TRSs tend to be A- and U-rich. The tables and junction logos obtained for the other two biological replicates gave similar results, and are reported in Suppl. Table 3 and Suppl. Fig. 2. The strategy used to collect the putative TRSs and to plot the logo are described in Material and Methods.

**Figure 7:**
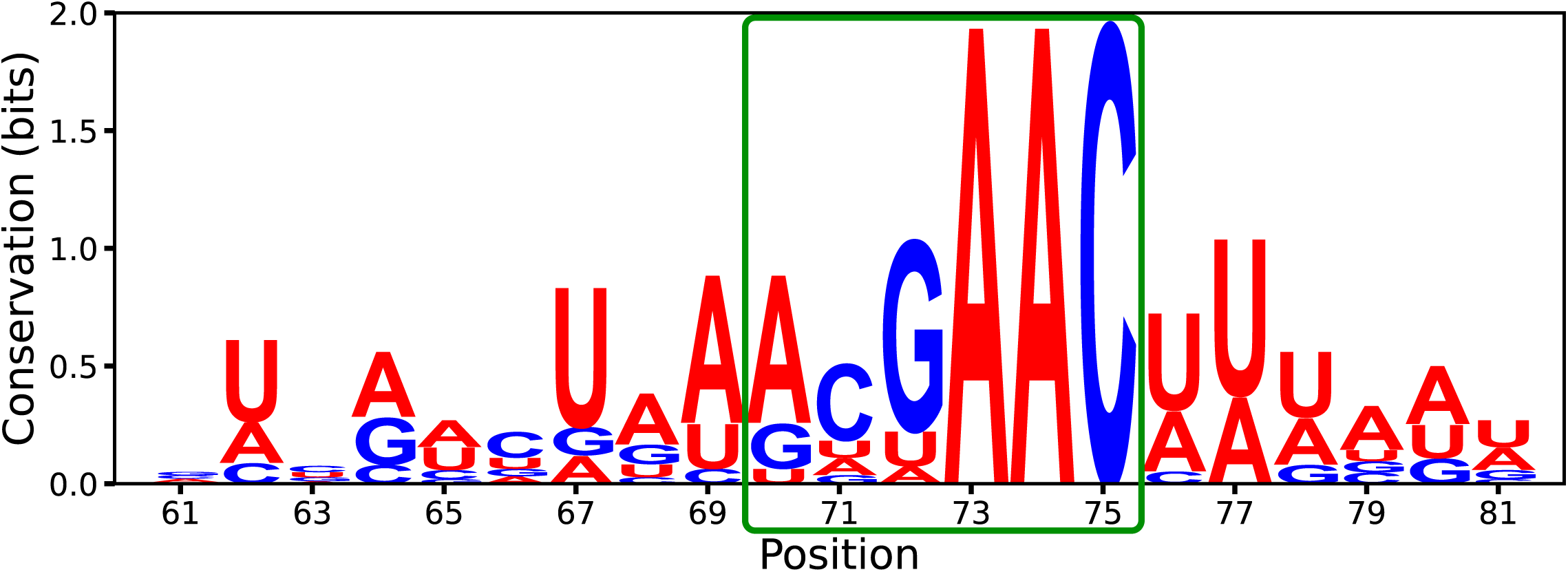
Logo of the RI positions around the TRS-putative sequences obtained from DI-tector. The conservation plotted as total height of the letters representing nucleotides is obtained as log_2_(4) *−∑_n_ f_i_*(*n*) log_2_(*f_i_*(*n*)), where *f_i_*(*n*) is the frequency of nucleotide *n* in position *i*. Therefore a height equal to 2 corresponds to perfect conservation. The horizontal axis are the position with respect to the reference sequence (GISAID ID: EPI ISL 414631). The green box highlights the canonical TRS. The alignment step to obtain this logo is described in the Material and Methods section. Color code used: red for adenine and uracil, blue for cytosine and guanine.

**Table 4:**
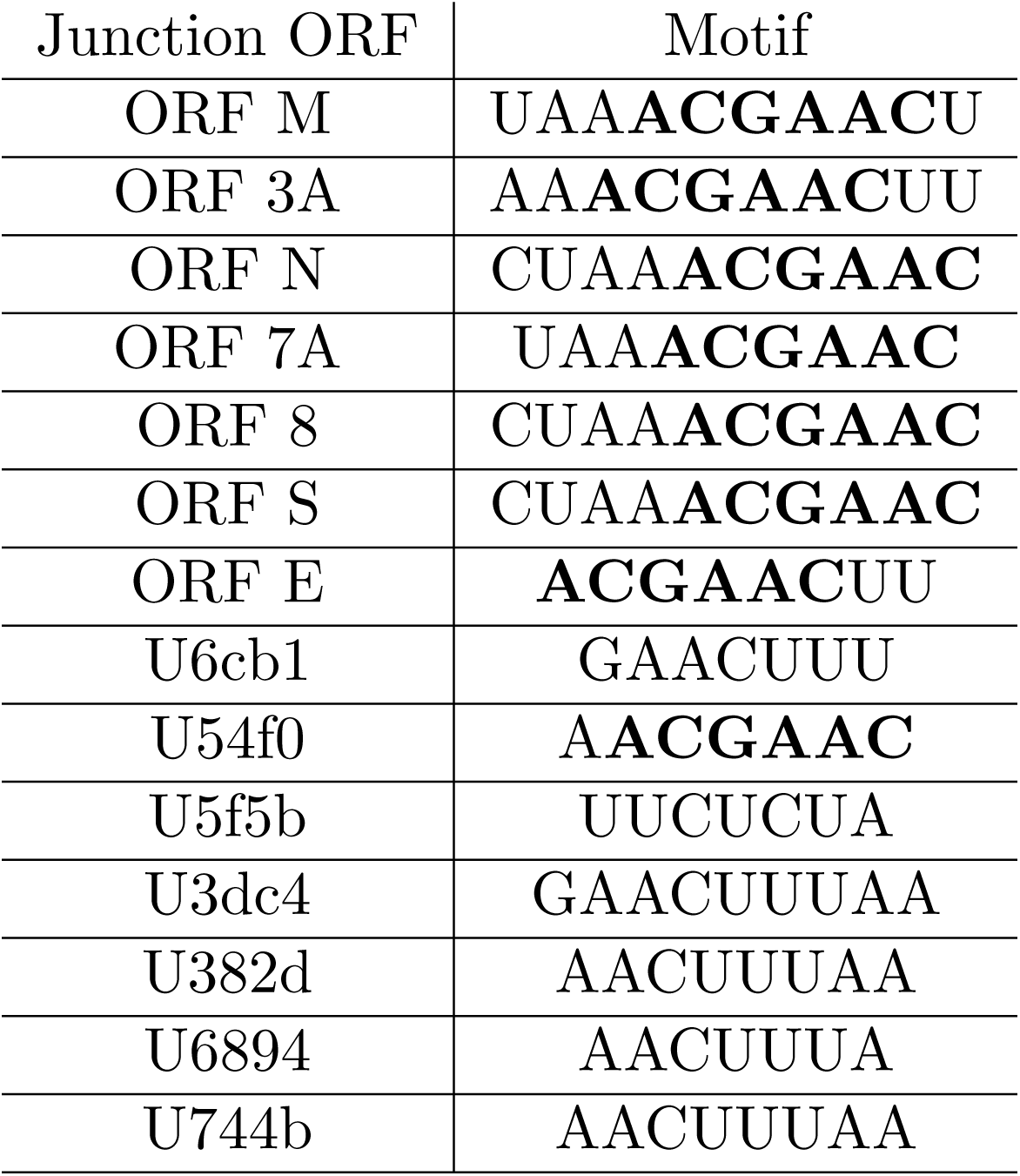
TRS and putative TRS detected by sgDI-tector. For SARS-CoV-2, sgDI-tector fixes the minimum length to have a TRS hit to 7 (see Material and Methods), so only junctions having a subsequence identical in the leader of length larger or equal 7 to are reported here. We highlighted the well-conserved canonical hexanucleotide ACGAAC motif with bold font. The results shown here have been obtained from the data coming from one biological replicate. The tables for the two additional biological replicates are given as Suppl. Table 3.

## Discussion

Based on suggested mechanisms of production for coronavirus sgRNAs [Pasternak et al., 2006, Sola et al., 2015, Wang et al., 2021], several bioinformatic approaches can be applied to detect them in NGS data: decumulation, detection of exon-exon junction reads (such as the STAR algorithm), alignment to leader sequence and TRS (such as the Periscope tool) and DVG recognition tools. We earlier developed DI-tector to characterize different forms of DVGs in NGS data. In this study we compared the other existing techniques for estimation and characterization of SARS-CoV-2 sgRNAs protocols with a pipeline based on the DI-tector’s output, which we named sgDI-tector.

A well-known strategy to quantify sgRNAs in coronaviruses is the so-called decumulation procedure, which has been introduced to characterize the sgRNA expression in cells infected by Murine coronavirus (also known as mouse hepatitis virus) [Irigoyen et al., 2016], and has been shown to give, in that case, consistent results with another standard approach consisting in finding the chimeric sequences that span the TRS. However, the correlation between the two approaches decreased when the results obtained by Finkel *et al*. on SARS-CoV-2-infected cells were analyzed. At the point that several sgRNA had negative RPKM after the decumulation procedure, highlighting another drawback of this method. Moreover, when the decumulation procedure was applied to our data, the results were completely inconsistent with any of the other tools used. We suggest that the low performance of the decumulation method for our data may be due to the large number of DVG observed, in particular insertions and deletions not resulting in canonical ORFs. This very large family of transcripts should be, in principle, accounted for in the decumulation procedure. Moreover, as observed in [Kim et al., 2020, Finkel et al., 2021], the canonical ORFs seem to be produced together with other, non-canonical ORFs and DVGs that should be considered also during decumulation. The remarkable correlation observed by Irigoyen *et al*. [Irigoyen et al., 2016] could be explained by the fact that most of DVGs and non-canonical ORFs are produced in relevant amounts later in infection, as was also discussed in [Finkel et al., 2021]. This was confirmed when DI-tector was run on Finkel *et al*.’s data: about 65% of the DVG at 5 hours post infection were associated with a canonical ORF and this number reduced to 46% at 24 hours post infection. Moreover, the total number of detected DVG (including canonical ORF junctions) compared with the number of non-DVG, mapped viral reads increased from 1% (5 hpi) to 2% (24 hpi).

Applying the STAR algorithm for detection of SARS-CoV-2 sgRNAs as a type of “exon-exon junction” was successfully performed in [Kim et al., 2020, Finkel et al., 2021, Wang et al., 2021]. We observed a strong correlation between NGS data analyzed by sgDI-tector and STAR algorithms with only few relevant differences. On the contrary, Periscope analysis on short-read data gave qualitatively different results, suggesting that this tool may have a decrease in performance when applied to a dataset not obtained through the ARTIC sequencing protocol for which the tool has been developed, especially when the reads have short lengths. When compared to previously published bioinformatic tools, the advantages of sgDI-tector are mainly two: firstly, sgDI-tector does not need to have as an input the leader sequence or the TRS, differently from Periscope and from other TRS-based approaches; secondly, sgDI-tector has been designed specifically for addressing the sgRNA level expression in viruses and for this reason it works without the need for unconventional parameter choices, as it is the case for STAR. In addition to these two technical advantages, sgDI-tector is user-friendly. We consider sgDI-tector to be the most user-friendly bioinformatic tool to estimate SARS-CoV-2 sgRNA from NGS data. Indeed, although STAR is a well-known mapping tool widely used in transcriptomics, it requires a large number of non-trivial options and parameters, which must be tuned accurately to detect sgRNAs. Periscope is a pipeline based on a workflow management system (snakemake) that is easy to install but, similarly to STAR, it has several parameters that the user must know and the full list of mandatory options is not provided. sgDI-tector is a Python script that is easy to run. It only needs the DI-tector script in the working directory (together with the tools required to run DI-tector, that is BWA and samtools), and all efforts have been made so that sgDI-tector operates with the lowest possible number of settings that can be easily selected based on virology knowledge. Finally, although the differences between STAR and sgDI-tector were small in most of the cases, for one canonical junction (used to express ORF 3A) the results of these two tools were statistically different, whereas Periscope’s result was for ORF 3A compatible with sgDI-tector’s result and not compatible with STAR’s result, suggesting that sgDI-tector might be more accurate than STAR in some cases (Table 3).

We showed here that the large fraction of deletion DVG detected by DI-tector can be used to identify and quantify viral sgRNAs from NGS data. Notice that, although all these ORFs have in principle the potential to express an ORF (5’ and 3’ identical to the full viral genome and an AUG start codon), we do not expect all of them to be translated, as their AUG codons could be within a poor Kozak context to serve as translation sites. Another possibility is that for some non-canonical ORFs the initiation of translation could be driven by non-AUG start codons [Kearse and Wilusz, 2017] and thus escape the sgDI-tector algorithm. Moreover, from our analysis we had access to the full set of DVG produced by SARS-CoV-2 during its life cycle, see Table 1. We used RT-qPCR to confirm the presence of detected by sgDI-tector non-canonical sgRNAs in cells infected with SARS-CoV-2. ORF U3dc4 is the non-canonical sgRNA that has its body TRS located in the ORF1ab and was detected in our three biological replicates specifically from infected cells (Fig. 3 and 4). This non-canonical RNA caught our attention as if transcribed it should encode a part of NSP12 protein which is SARS-CoV-2 RdRp. However additional experiments are required to validate that ORF U3dc4 is indeed translated.

In addition to the deletion DVGs, we also detected an important number of insertion DVGs. sgDI-tector algorithm can only detect insertions of viral origin. Notice that the observed number of insertion DVG reads (average over replicates: 23.8% of all the DVG reads) is very close to the number of deletion DVG reads that cannot be associated with canonical sgRNAs (average over replicates: 23.4% of all the DVG reads). Further studies are needed to understand whether these insertion events correspond to the real DVGs produced during viral replication or represent viral genome recombination events previously described for coronaviruses [Simon-Loriere and Holmes, 2011].

Finally, we detected a very low number of DVG of the type 5’/3’-copy-backs/snap-backs. Of note, 5’/3’-copy-back DVG are largely described for negative-sense RNA viruses [Lazzarini et al., 1981, Dimmock et al., 2014].

Exact mechanisms of production of DVG are unknown. The central question is whether their production is induced by host factors, aiming at introducing interferences with viral replication and allowing virus detection by the host’s innate immune system. As stated, DVG are truncated and/or rearranged forms of viral genomes generated by most viruses during viral replication and sharing the minimum essential characteristics for replication: a competent initiation site at the 3’-end, its complementary sequence at the 5’-end. It is intriguing that the above description can also be applied on SARS-CoV-2 sgRNAs. This similarity can further be used to suggest mechanisms for production of deletion forms of DVG arguing for an internal property of viral RdRp to produce DVGs. Thus, further comparison of molecular structures and kinetics of coronavirus sgRNAs and DVGs accumulation will be of strong interest for future studies.

## Materials and Methods

### Cells

HEK293 (human embryonic kidney cells) cell lines expressing One-STrEP-tagged Cherry (ST-CH) [Sanchez David et al., 2016], Vero-E6 (African green monkey kidney cells, ATCC CRL-1586), were maintained at 37°C, 5% CO2 in Dulbecco’s modified Eagle medium (DMEM; Thermo Fisher Scientific) supplemented with 5% heat-inactivated fetal calf serum (FCS; GE Healthcare) and 1% PS (Penicillin 10,000 U/mL; Streptomycin 10,000 *µ*g/mL). ST-CH cell line was supplemented with G418 (Sigma) at 500 g/ml. The absence of mycoplasma was regularly checked by RT-PCR in all cell lines. For the generation of ST-CH overexpressing ACE-2 (ST-CH^ACE-2^) lentivirus transduction of hACE2 was performed. Cells were screened by FACS for ACE2 expression and susceptibility to SARS-CoV-2 replication.

### Viral titers, infection with SARS-CoV-2 and total RNA extraction

The SARS-CoV-2 hCoV-19/France/GES-1973/2020 GISAID ID: EPI ISL 414631 strain was supplied by the National Reference Centre for Respiratory Viruses hosted by Institut Pasteur (Paris, France). Vero-E6 cells were used for the amplification and titration of viral stocks. Vero-E6 monolayers were infected with SARS-CoV-2 in the presence of 0.1% TPCK trypsin (Sigma) at a multiplicity of infection (MOI) of 0.0001 plaque-forming units (pfu) per cell. When the cytopathogenic effect was apparent, the culture supernatant was collected and centrifuged for 5 minutes at 850g. The efficiency of virus amplification was evaluated by titrating the supernatant on Vero-E6 cells, in a standard plaque assay adapted from Matrosovich *et al*. [Matrosovich et al., 2006]. For SARS-CoV-2 infection ST-CH^ACE-2^ cells were seeded into polylysine-coated (SIGMA) T150 flasks 1 day before infection (20×106 cells/flask). Virus infections were carried out at an MOI of 1. Viruses were diluted with DMEM 0%FCS to obtain a final inoculum volume of 5 ml. Cells were incubated with virus for 1 h at 37°C with gentle shaking. Twenty-five milliliters of DMEM containing 0%FCS was added to each T150 flask, and cells were incubated at 37°C until infections were stopped by cell lysis 24 h later. Total RNA was extracted from either SARS-CoV-2- or mock-infected ST-CH^ACE-2^ using TriLS (TriLS, Sigma) reagent protocol previously described in details in [Sanchez David et al., 2016]. All experiments with SARS-CoV-2 were conducted under strict BSL3 conditions.

### Raw data collection, pre-processing and normalization scheme

NGS libraries were built using a TruSeq mRNA-Seq library preparation kit (Cat#20020594 Illumina), according to the manufacturer’s recommendations. Quality control was performed on an Agilent Bioanalyzer. Sequencing was performed on the Illumina NextSeq500 platform to generate single-end 75 bp reads bearing strand specificity. Reads were cleaned of adapter sequences and low-quality sequences using cutadapt version 2.9. Only sequences at least 25 nucleotides in length were considered for further analysis. Bowtie version 2.1.0, with default parameters, was used for alignment on the reference genome (hCoV-19/France/GES-1973/2020, GISAID accession Id: EPI ISL 414631). SARS-CoV-2 genome coverages were computed with bedtools genomecov for each strand.

For several analyses performed in this work we needed to use data coming from the three biological replicates of our experimental setup. To compare in a more robust way the data coming from independent experiments, we normalized all the counts as follows: for each biological replicate, we took the number of reads mapped to the SARS-CoV-2 genome and divided this by the value obtained for Replicate 1 (the raw number of reads are reported in Suppl. Table 1). The three values that we obtained in this way (1 for Replicate 1, and other values for other replicates) are used to re-scale all the number of reads before taking any average over biological replicates. In particular, averages of normalized data have been used for Table 3 and 1, and for Figs. 2 and 4.

### RT-qPCR validation of non-canonical U3dc4 ORF

RT-qPCR was performed on RNA samples from SARS-CoV-2- or mock-infected cells performed in three biological replicates and prepared as described for NGS. First-strand complementary DNA (cDNA) synthesis was performed on 2500 ng of total RNA in a final volume of 20 µL with the Superscript IV vilo (Thermo Scientific #11756050) according to the manufacturer’s protocol. RT-qPCR analysis was performed using Applied Biosystems StepOnePlus technology. Reactions were performed on an equivalent of 50ng of total RNA by use of SYBER Green Kit (Thermo Fisher Scientific #4309155) for qPCR analyze according to the manufacturer’s protocol. Reactions were performed in a final volume of 20 µL in the presence of 60 nM U3dc4-specific forward (5’-CCTTCCCAGGTAACAAAC) and reverse (5’-GTCTCAGTCCAACATTTTG) primers; or N-specific forward (5’-TAAAGGTTTATACCTTCCCA) or reverse (5’-CGTTCTCCATTCTG-GTTA) primers; or GAPDH forward (5’-CACATGGCCTCCAAGGAGTAA) and reverse (5’-TGAGGGTCTCTCTCTTCCTCTTGT) primers.

### sgDI-tector pipeline: from NGS data to sgRNA detection

When sgDI-tector is run, it firstly calls DI-tector [Beauclair et al., 2018] (here used in version 0.6) with default parameters (using bwa v0.7.17, bedtools v2.17.0 and samtools v1.9) to detect SARS-CoV-2 DVGs. DI-tector outputs four different types of DVG, namely deletions, insertions, 3’- and 5’-copy backs/snapbacks. For each deletion, the BP (breakpoint) and RI (reinitiation) site are specified, consisting in the two sites that, despite being separated in the full-length genome, are brought together in the junction read. The pipeline for sgRNA detection starts by finding the window of a user-modifiable length (the default value is 20) with the largest number of BP. Under the hypothesis that the virus is replicating, and that replication needs sgRNAs, we expect (and verify in each *in vitro* sample analyzed here) that this window coincides with the end of the leader sequence. Then, sgDI-tector filter deletion DVGs by requiring the BP to lay into this window. The resulting deletions are then associated with an ORF, by finding the first ATG sub-sequence after the RI. The families of junctions obtained in this way are then sorted by the number of reads belonging to the family, and given as an output. Optionally the user can provide a list of reference subgenomic ORFs (in fasta format), that will be used by sgDI-tector to name the sgRNAs found. In particular, this is done by aligning the putative expressed protein to the list of known proteins, and comparing the alignment score with a fixed threshold. When more than one hit is obtained for the same viral protein (for instance, because two different sgRNA produce two proteins with few amino acid of difference), the name of the top hit is take from the user-provided file, and the name of successive hits are obtained from it by adding increasing numbers (e. g. ORF 3A-1). In addition to the sgRNA list, sgDI-tector outputs the leader sequence used. The full sgDI-tector pipeline described here, which takes as input the DI-tector results and gives as output a list of putative sgRNAs, is presented in Fig. 8. In all samples analyzed, all the observed canonical subgenomic ORFs (S, 3A, E, M, 6, 7a, 7b, 8, N) were within the 13 families with the largest number of reads (see Fig. 3 and Fig. 4).

**Figure 8:**
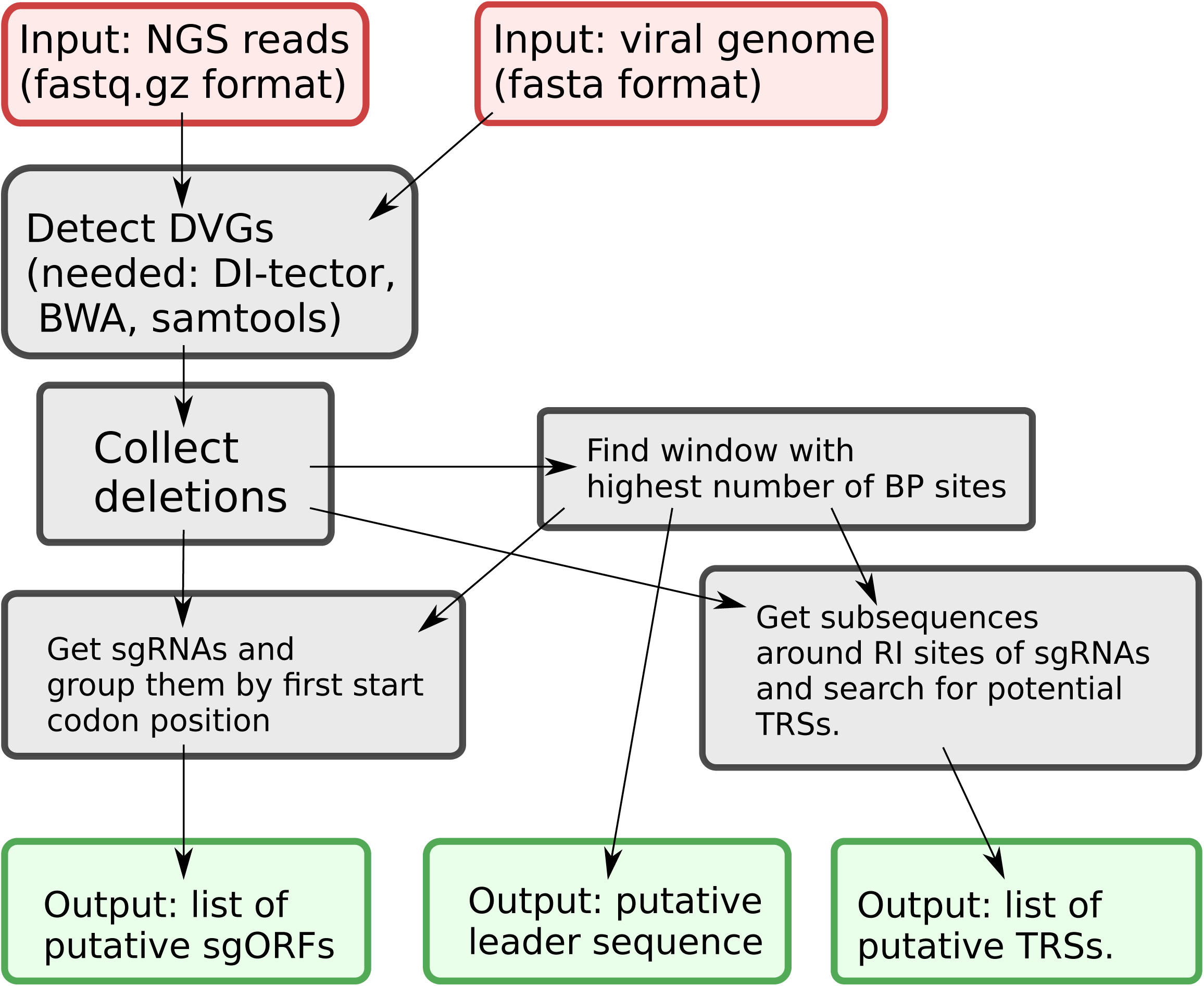
Scheme of the sgDI-tector pipeline. introduced here to find the putative position of the leader sequence, sgRNAs, and a list of putative transcription-regulatory sequences (TRSs). Red boxes denotes necessary inputs for the sgDI-tector tool, and green boxes denotes outputs.

### TRS detection through junction analysis

DI-tector itself allows several graphical outputs, and among them there is the sequence logo of nucleotides just before and after the RI position. However, this functionality cannot be used as it is for sgRNA junctions. Indeed, it is well known [Sola et al., 2015] that the leader-body junction is regulated by a core subsequence that is identical in the two sides of the junction, and this creates ambiguity in defining precisely BP and RI, and makes a further alignment step necessary to correctly compare short subsequences spanning the RI position. However, the alignment step is non-trivial, as these subsequences are typically not alignable but for some small parts. Therefore, we used a different approach: firstly, sgDI-tector computes the probability of a random sub-sequence of the viral genome to have a sub-sequence of length *L* (putative TRS) which appears also in the final part of the leader sequence. This allows sgDI-tector to fix *L^∗^*, the length for which this probability is lower than 0.05. Then, for all sequences spanning the RI position in junctions, putative TRS of length *≥ L^∗^* are collected, and saved in an output file. Once the putative TRSs have been obtained, they can be aligned to the leader sequence. The resulting logo, which has been directly obtained from the sequences in Table 4, is presented in Suppl. Fig. 3. To have an even more complete information about the body side of the junctions, once the putative TRSs are aligned, the remaining nucleotides of the body side of the junctions can be used together with TRSs to produce the logo presented in Fig. 7 (and Suppl. Fig. 2 for the other two biological replicates). Equivalently, the logos are obtained from the sequences around the body part of the junctions listed in Table 4, aligned to the leader sequence so that the TRSs obtained in Table 4 overlap with the corresponding identical sequence in the leader part. Finally, we decided not to include any information about abundances of the junctions in the logos. If such an information would be included, of course, the full canonical TRS 5’-ACGAAC-3’ (highlighted by a green box in Fig. 7) would become almost perfectly conserved, because it is present in all the junctions with high number of counts.

## Supporting information

Suppementary information file.

## Acknowledgments

We would like to thank J. Pipoli da Fonseca, L. Lemée from Biomics Platform, C2RT, Institut Pasteur, Paris, France for RNA NGS, IBISA and the Illumina COVID-19 Projects’ offer. The authors would like to thank members of the Tangy, van der Werf laboratories for support and valuable discussions. We acknowledge the ANR (Agence Nationale de la Recherche) and FRM (Fondation de la Recherche Médicale) for funding this work through the AAP Flash-Covid 19 project SARS-Cov-2immunRNAs. AVK received support from ANR through the grant ANR-LBX-62 IBEID CoV-2SENSING/COVID 19. RNA NGS has been supported by France Génomique (ANR-10-INBS-09-09).

## Availability of data and materials

The data collected and used for this work have been deposited in NCBI’s Gene Expression Omnibus [Edgar et al., 2002] and are accessible through GEO Series accession number GSE180632, at https://www.ncbi.nlm.nih.gov/geo/query/acc.cgi?acc=GSE180632. sgDI-tector code is publicly available on Github, at https://github.com/adigioacchino/sgDI-tector.

