## Supplementary material for "sgDI-tector: defective interfering viral genome bioinformatics for detection of coronavirus subgenomic RNAs": Suppementary information file.

### Supplementary Information

**Andrea Di Gioacchino, Rachel Legendre, Yannis Rahou, Valérie Najburg, Pierre Charneau, Benjamin D. Greenbaum, Frédéric Tangy, Sylvie van der Werf, Simona Cocco, Anastasia V. Komarova**

| | Replicate 1 | Replicate 2 | Replicate 3 |
|---|---|---|---|
| Total number of reads | 34470773 | 39467941 | 42751723 |
| Human genome<br>(% of total reads) | 18351424<br>(53.2%) | 18855674<br>(47.7%) | 20219414<br>(47.3%) |
| Human ribosomal RNA<br>(% of total reads) | 119155<br>(0.3%) | 115706<br>(0.3%) | 156747<br>(0.4%) |
| Viral (SARS-CoV-2)<br>(% of total reads) | 11461575<br>(33.2%) | 15714257<br>(39.8%) | 16776050<br>(39.2%) |
| Total DVG reads<br>(% of viral reads) | 81866<br>(0.7%) | 80598<br>(0.5%) | 94319<br>(0.6%) |
| deletion DVG<br>(% of total DVG) | 59886<br>(73.2%) | 60854<br>(75.5%) | 70133<br>(74.4%) |
| of which sgRNA<br>(% of deletion DVG) | 41751<br>(69.7%) | 41029<br>(67.4%) | 47858<br>(68.2%) |
| insertion DVG<br>(% of total DVG) | 20577<br>(25.1%) | 18011<br>(22.3%) | 22409<br>(23.8%) |
| copyback DVG<br>(% of total DVG) | 1403<br>(1.7%) | 1733<br>(2.2%) | 1777<br>(1.9%) |

Supplementary Table 1: Summary of results of RNAseq and alignment of the reads to the human genome, and SARS-CoV-2 viral genome. NGS library preparation was performed with an rRNA depletion step. The unmapped reads have been further processed with DI-tector and the resulting characterization of the DVG reads into deletions, insertions and copy-backs is given after the double horizontal line. To count sgRNA reads, we only considered standard subgenomic ORFs for SARS-CoV-2 (S, 3A, E, M, 6, 7a, 7b, N, 10). The DVG copyback row contains reads coming from both 3' and 5' copybacks and snapbacks.

| ORF name | AUG start position in genome |
|---|---|
| U6cb1 | 27825 |
| U54f0 | 21744 |
| U633d | 25405 |
| U59dc | 23004 |
| U5345 | 21317 |
| U7226 | 29222 |
| U5757 | 22359 |
| U5856 | 22614 |
| U3dc4 | 15812 |
| U11ee | 4590 |
| U5df3 | 24051 |

Supplementary Table 2: Conversion between names of the non-standard ORFs with highest counts obtained by sgDI-tector (blue bars in Figs. 4 and 5) and the corresponding position of the start codon (AUG) in the SARS-CoV-2 genome. The conversion is made by interpreting the alphanumerical characters after the U (which stands for unknown) in the ORF name as an hexadecimal number. The position in genome is given with respect to the SARS-CoV-2 strain hCoV-19/France/GES-1973/2020, with GISAID ID: EPI_ISL_414631.

| Replicate #2 | | Replicate #3 | |
|---|---|---|---|
| Junction ORF | Motif | Junction ORF | Motif |
| ORF M | UAA**ACGAAC**U | ORF M | UAA**ACGAAC**U |
| ORF 3A | AA**ACGAAC**UU | ORF 3A | AA**ACGAAC**UU |
| ORF N | CUAA**ACGAAC** | ORF N | CUAA**ACGAAC** |
| ORF 7A | UAA**ACGAAC** | ORF 7A | UAA**ACGAAC** |
| ORF 8 | CUAA**ACGAAC** | ORF 8 | CUAA**ACGAAC** |
| ORF S | CUAA**ACGAAC** | ORF S | CUAA**ACGAAC** |
| U6cb1 | GAACUUU | ORF E | **ACGAAC**UU |
| ORF E | **ACGAAC**UU | U6cb1 | GAACUUU |
| U54f0 | A**ACGAAC** | U54f0 | A**ACGAAC** |
| ORF 3A-1 | A**ACGAAC**UU | ORF 3A-1 | A**ACGAAC**UU |
| U5757 | AACUUUA | U3dc4 | GAACUUUAA |
| U3dc4 | GAACUUUAA | U5757 | AACUUUA |
| | | U5f5b | UUCUCUA |
| | | U6d2e | UAA**ACGAAC** |
| | | U744b | AACUUUAA |
| | | U52c4 | CUCUAAA |

Supplementary Table 3: TRS and putative TRS obtained for the two biological replicates not discussed in the main text. The double vertical line divides the two biological replicates. The junction ORFs are sorted according to the number of times that junction was found by the algorithm (in decreasing order). As it is apparent, the results are consistent across replicates, especially for the junctions with more counts.

| | Replicate #2 | | Replicate #3 | |
|---|---|---|---|---|
| Target detected | SARS-CoV-2 | Non-infected | SARS-CoV-2 | Non-infected |
| U3dc4 | $23.72 \pm 0.04$ | ND | $22.1 \pm 0.1$ | ND |
| ORF N | $13.9 \pm 0.1$ | ND | $12.6 \pm 0.1$ | ND |
| GAPDH | $19.80 \pm 0.03$ | $18.8 \pm 0.1$ | $19.5 \pm 0.1$ | $18.5 \pm 0.1$ |

Supplementary Table 4: **RT-qPCR validation of U3dc4 sgRNA transcript.** RT-qPCR $C_t$ values for U3dc4, ORF N, and GAPDH detection in cDNA equivalent of 50 ng of total RNA extracted from SARS-CoV-2 or mock-infected HEK293T cells are shown. Samples were analyzed in duplicates. ND: Not Determined ($C_t > 34$). Data are given as average $\pm$ standard deviation. Results obtained from the two biological replicates not shown in Table 2.

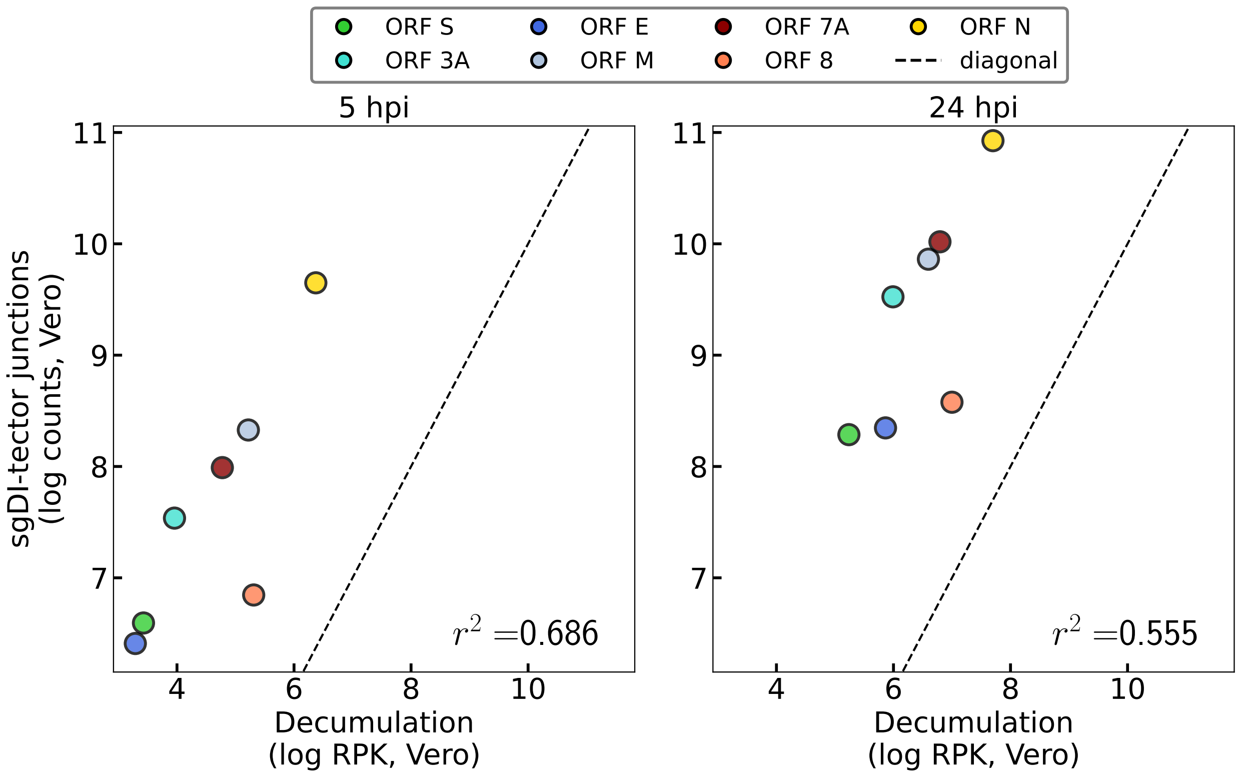


Supplementary Figure 1: sgDI-tector detected junctions are more correlated with the results of the decumulation method when the two strategies are applied to Finkel *et al.* 's data. In particular, this correlation is higher at 5 hpi, when less DVG are present.

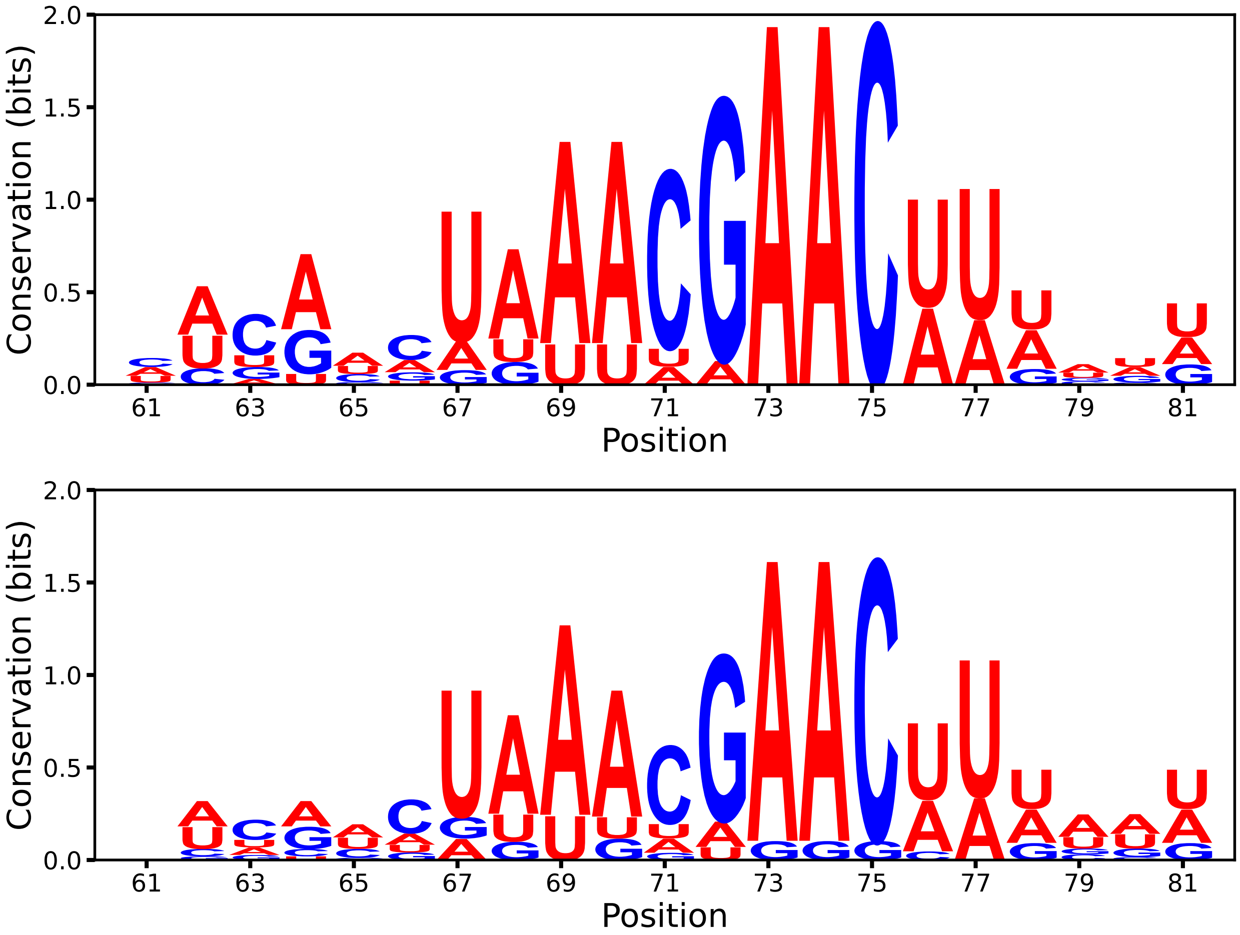


Supplementary Figure 2: Sequence logo around RI positions aligned to the leader sequence, for the two other biological replicates (see Fig. 7).

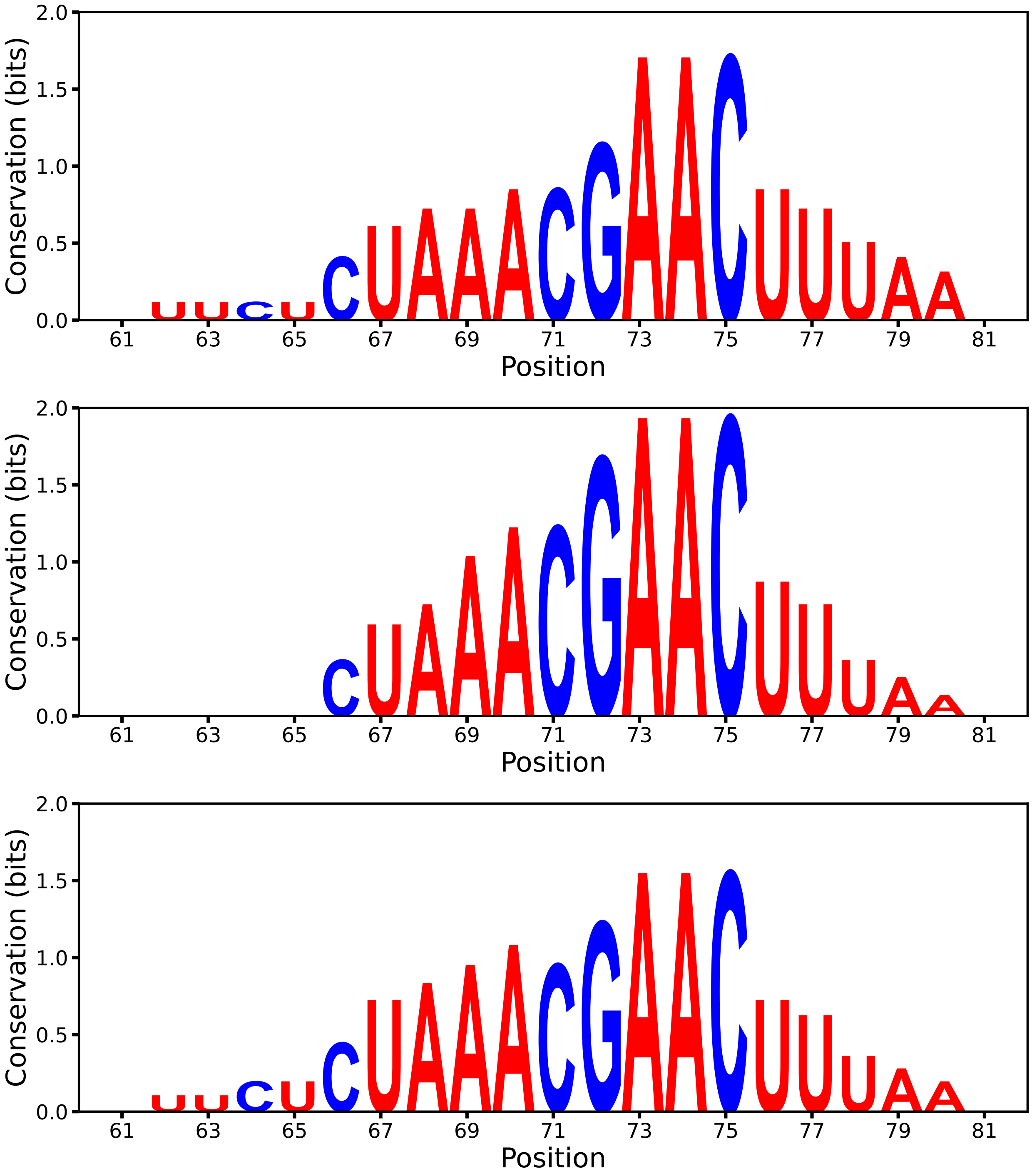


Supplementary Figure 3: Sequence logo for the putative TRS sequences only, for the three biological replicates (see Tab. 2).
